# Two intramolecular interfaces control the WDR44 conformational switch that regulates Rab11 binding

**DOI:** 10.64898/2026.09.10.750756

**Authors:** Hailey Eng, Michael Davey, Elizabeth Conibear

## Abstract

Rab GTPases coordinate distinct membrane trafficking pathways through interactions with specialized effector proteins. Rab11 is a central regulator of endocytic recycling and also drives preciliary trafficking to the centriole. Both pathways depend on the Rab11 effector WDR44, which promotes Rab11-dependent recycling but suppresses ciliogenesis in serum-grown cells. WDR44 engages Rab11 through an N-terminal Rab11-binding domain that is autoinhibited by an intramolecular interaction with the C-terminal WD40 domain, and pathogenic WD40-domain variants that cause ciliopathies relieve this inhibition. How the closed conformation is maintained, however, has not been defined. Here we combine structural modeling and mutational analysis with a bystander BRET assay that reports WDR44 recruitment to Rab11 compartments in living cells, and identify two N-terminal segments that engage distinct surfaces of the WD40 domain. The larger segment extends across the top face of the domain, where reported pathogenic variants cluster, while a second segment contacts the side. Disrupting either segment weakens the intramolecular interaction and promotes WDR44 recruitment to Rab11 compartments. Our work defines the contacts that maintain WDR44 in its closed conformation and provides a mechanism by which pathogenic mutations may alter WDR44 function.

**Significance statement:**

- Rab11 cooperates with effector proteins, including WDR44, to coordinate intracellular trafficking during endosomal recycling and ciliogenesis. A closed conformation of WDR44 blocks Rab11-binding and is released by disease-causing mutations, but intramolecular contacts that maintain autoinhibition have not been identified.
- Using structural modeling and cell-based analyses, we identify two regions of WDR44 that fold onto distinct surfaces of its own C-terminal domain, and can be released to promote Rab11 association.
- These contacts explain how disease-causing mutations activate WDR44, and define the surfaces through which its functions are likely regulated.

## Introduction

Delivery of intracellular traffic to specific compartments is regulated by Rab GTPases, which coordinate distinct trafficking pathways through interactions with their effector proteins when in a GTP-bound state. Rab11 is a central regulator of endosomal recycling to the plasma membrane (PM) and is primarily found at recycling compartments (Ullrich *et al*., 1996, Lock and Stow, 2005). These compartments can be heterogenous in both composition and morphology but are often described as dynamic tubular endosomes, or the tubular endosome network (TEN) (Sönnichsen et al., 2000, Goldenring, 2015, Farmer et al., 2021). In addition to its role in recycling, Rab11 promotes ciliogenesis by initiating a Rab cascade that enables preciliary trafficking (Knödler *et al*., 2010, Westlake *et al*., 2011).

Among Rab11 effectors, WDR44 (also named Rab11BP/Rabphilin-11; Zeng *et al*., 1999, Mammoto *et al*., 1999) has emerged as a key regulator of both recycling and ciliogenesis. WDR44 is required for the PM-directed recycling and export of various cargoes including E-cadherin, MMP14, and CFTR (Lucken-Ardjomande Häsler *et al*., 2020). It is enriched at a subset of recycling endosomal tubules that form membrane contacts with the endoplasmic reticulum (ER). At endosomes, WDR44 associates with the tubulating BAR-domain protein GRAF1/2 via a proline-rich sequence, while it anchors to the ER through a FFAT motif that binds VAPA/B (Baron et al., 2014, Lucken-Ardjomande Häsler *et al*., 2020, Parolek and Burd, 2024).

More recently, WDR44 was proposed to inhibit ciliogenesis in serum-grown cells by competing for Rab11 binding to prevent formation of the Rab11-Rabin8-FIP3 complex required for Rab8 activation and delivery to the centriole (Walia *et al*., 2019). The cytosolic pool of WDR44 exists in a closed conformation where the Rab11-binding domain, in its largely unstructured N-terminus, is inhibited by interactions with a WD40 repeat domain in the C-terminus (Zeng *et al*., 1999, Mammoto *et al*., 1999, Thibodeau *et al*., 2023). When this C-terminal domain is deleted, the N-terminal fragment binds Rab11 and redistributes to Rab11-positive structures, typically enlarged endosomes at the cell periphery (Lucken-Ardjomande Häsler *et al*., 2020).

A similar increase in Rab11 binding is caused by pathogenic variants of WDR44 that cause ciliopathies, developmental disorders resulting from defective cilia (Accogli *et al*., 2024). These pathogenic variants map to the top face of the WD40 domain, and decrease binding between N- and C-terminal fragments in vitro, suggesting they are gain-of-function mutations that disrupt the WDR44 autoinhibitory conformation. However, the specific interfaces that maintain this autoinhibitory conformation have not been defined.

Here, we investigated the basis of WDR44 autoinhibition that regulates WDR44-Rab11 complex formation. We established a sensitive bystander BRET assay to detect changes in WDR44-Rab11 proximity in the cellular context. Using in silico modeling, we identified two main regions in the WDR44 N-terminus predicted to contact the C-terminal WD40 domain. Mutational analysis shows that both interfaces contribute to intramolecular interactions and that their disruption is sufficient to induce Rab11 association. Together, our findings detail the intramolecular interfaces that control WDR44 conformation, and implicate these contacts as candidate sites of regulation.

## Results and Discussion

### Bystander BRET detects recruitment of pathogenic WDR44 to Rab11 compartments

To identify the mechanism underlying autoinhibition, we took advantage of the observation that the open conformation enhances Rab11 binding and redistributes WDR44 from the TEN to Rab11-labeled structures (Zeng *et al*., 1999, Lucken-Ardjomande Häsler *et al*., 2020). Transiently expressed WDR44 is found on a heterogeneous set of tubules, small puncta, and larger structures (Parolek and Burd, 2024, Accogli et al., 2024, Lucken-Ardjomande Häsler et al., 2020), making it challenging to quantify localization using conventional fluorescence imaging. To measure the cellular distribution of Rab11-bound WDR44, we instead exploited a sensitive bystander bioluminescence resonance energy transfer (BRET) assay (Lan *et al.,* 2012, Holme, Sapia *et al*., 2025). A positive localization readout, or ‘net BRET’ signal, is produced when WDR44 tagged with the RLuc8 luciferase (donor) excites an abundant Venus-tagged organelle marker (acceptor) in close proximity (Figure 1A), providing a quantitative readout of localization changes with minimal background signal.

**Figure 1.**
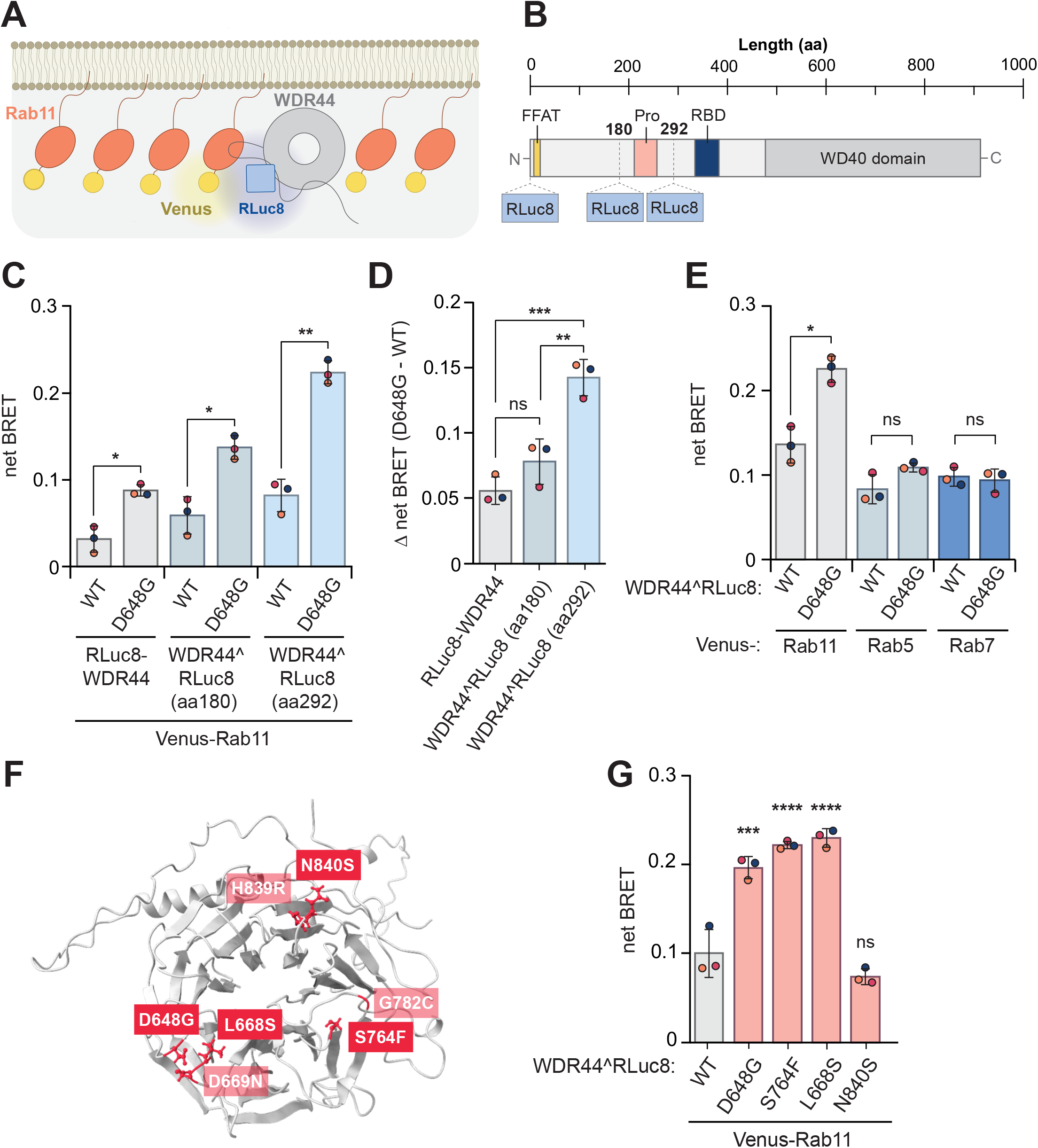
Bystander BRET detects recruitment of pathogenic WDR44 variants to Rab11 compartments. **(A)** Schematic of the bystander BRET assay used to detect organellar localization of a protein of interest. When RLuc8-tagged WDR44 (donor) is in close proximity to overexpressed Venus-tagged Rab11 (acceptor), a net BRET signal is produced, representing localization of WDR44 at Rab11 membranes. **(B)** Schematic of internal RLuc8 tag insertion sites within WDR44. **(C)** BRET ratios indicate WDR44^D648G^ is recruited to Rab11-positive compartments when it is tagged N-terminally or internally at two sites. HEK293T cells were transiently transfected with RLuc8-tagged WDR44, WT or D648G, and Venus-Rab11. Repeated measures two-way ANOVA with Holm-Šídák’s multiple comparison’s test; n = 3; ns = P > 0.05, * = P ≤ 0.05, ** = P ≤ 0.01. Error bars indicate standard deviation. **(D)** Differential co-localization of WT and D648G mutant protein with Rab11 is highest when they are internally tagged with RLuc8 between amino acids 292-293. Differences between WT and mutant protein recruitment from C were calculated and compared across tagging locations. One-way ANOVA with Tukey’s multiple comparison’s test; n = 3; ns = P > 0.05, ** = P ≤ 0.01, *** = P ≤ 0.001. Error bars indicate standard deviation. **(E)** Increased localization of pathogenic WDR44 is specific to Rab11 endosomes. Repeated measures two-way ANOVA with Holm-Šídák’s multiple comparison’s test; n = 3; ns = P > 0.05, * = P ≤ 0.05. Error bars indicate standard deviation. **(F)** Schematic showing disease-causing mutation sites which are enriched on the top surface of the WD40 domain. Mutation sites evaluated in this study are shown in brighter red. **(G)** BRET ratios indicate that the D648G, S764F, and L668S, but not N840S, disease variants are recruited to Rab11-positive compartments. One-way ANOVA with Dunnett’s multiple comparison’s test; n = 3; ns = P > 0.05, *** = P ≤ 0.001, **** = P < 0.0001. Error bars indicate standard deviation. BRET, bioluminescence resonance energy transfer; FFAT, two phenylalanines in an acidic tract; Pro, proline-rich motif; RBD, Rab11-binding domain.

To determine if recruitment of WDR44 to Rab11-positive compartments can be measured by bystander BRET, we compared the localization of the WT protein to a gain-of-function pathogenic mutant, D648G, tagged with the RLuc8 luciferase. Although previous studies used N-terminally tagged WDR44 (Lucken-Ardjomande Häsler et al., 2020, Accogli *et al*., 2024, Parolek and Burd, 2024), the N-terminus (residues 1-15) contains the ER-binding FFAT motif (Baron et al., 2014), placing the N-terminal tag far from endosomal Rab11, found on the opposing side of the organelle contact site. To improve detection of WDR44 at endosomes, we tested whether insertion of RLuc8 at internal sites would enhance its ability to excite Venus-Rab11. We identified two optimal sites in the unstructured region of the WDR44 N-terminus that were outside of any known binding domains in that region including the FFAT motif, proline-rich motif, and Rab11-binding domain (Figure 1B). By BRET analysis, the WT protein showed a low level of recruitment to Rab11-positive structures, whereas the D648G variant showed an increase in Rab11 proximity, consistent with previous reports (Accogli *et al*., 2024) (Figure 1C). Differential localization of the D648G mutant was greater when it was tagged at internal sites (WDR44^RLuc8(aa180) and WDR44^RLuc8(aa292)) and was optimally detected when RLuc8 was inserted between residues 292-293 (hereafter WDR44^RLuc8) (Figure 1, C and D). These results demonstrate that bystander BRET is a sensitive assay that recognizes changes in the intracellular distribution of WDR44 in complex with Rab11.

WDR44 enrichment at endosomes is specific to Rab11-labeled compartments, as the D648G variant showed no increased proximity to Venus-tagged Rab5 or Rab7 (Figure 1E). We next used BRET to measure the effect of three additional pathogenic mutations, each in a distinct region of the WD40 domain, which were previously reported to enhance Rab11 binding (Accogli *et al.,* 2024) (Figure 1F). The S764F, L668S and D648G variants showed a robust increase in Rab11 proximity, while the N840S mutant, which exhibits weaker Rab11 binding and was not previously observed at Rab11-positive compartments (Accogli et al., 2024), did not produce a significant BRET signal (Figure 1G). Therefore, bystander BRET can distinguish differential Rab11 recruitment across a range of WDR44 gain-of-function variants.

### Two segments of the WDR44 N-terminus contact distinct surfaces of the WD40 domain

The WDR44 C-terminus limits the binding of the N-terminus to Rab11, suggesting interactions between the two segments are autoinhibitory (Zeng *et al*., 1999), though the intramolecular interactions have not been defined. To identify inhibitory contacts in WDR44, we used AlphaFold2 (Jumper *et al.,* 2021) to model the N-terminal (1-504, WDR44^NT^) and C-terminal (477-913, WDR44^CT^) fragments as separate chains. A confident interaction was predicted between the WDR44 fragments, supported by a high interface predicted template modeling (ipTM) score (Figure 2A), and pLDDT values (Figure S1A).

**Figure 2.**
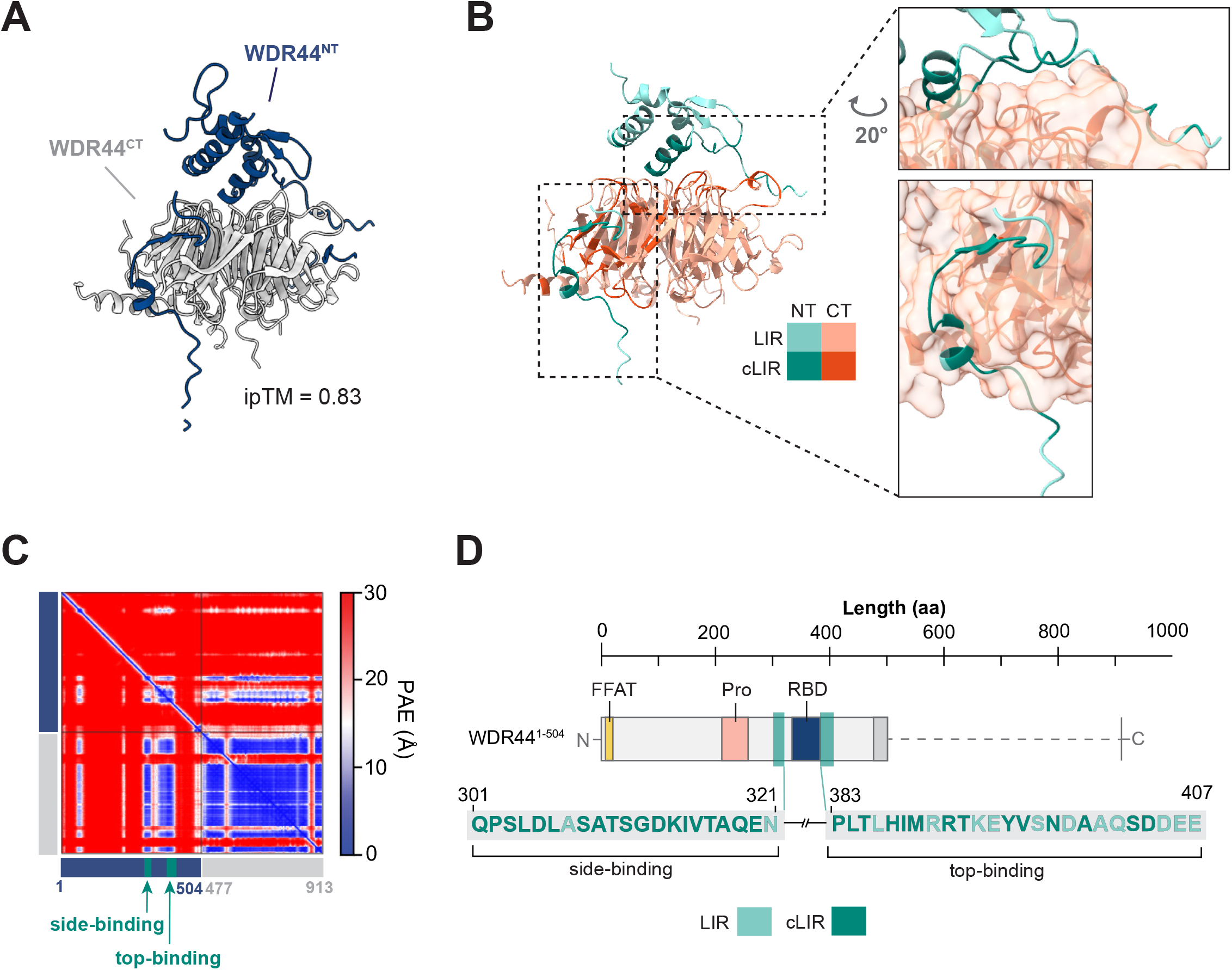
Structural predictions identify two interdomain binding interfaces**. (A**) AlphaFold2-predicted structure of the WDR44 N-terminus (blue) modeled against the C-terminus (grey). The highest ranked model is shown and residues with pLDDT values < 30 are hidden. ipTM is indicated. **(B)** Interactions between the WDR44 N- and C-termini are predicted at two interfaces along the top and side of the WD40 domain, as shown by LIVIA-computed local interaction residues (LIR, light green/orange) and the subset of these that are also in direct contact (cLIR, dark green/orange), based on the AlphaFold2-predicted structure shown in A. **(C)** PAE plot for the highest-ranked model of the AlphaFold2-based predictions shown in A. The regions with the highest confidence, where the side-binding segment and top-binding helix interface with the WDR44 C-terminus, are indicated by arrows. **(D)** Schematic of WDR44 N- and C-terminal fragments and intramolecular interfaces, with LIVIA-computed LIR (light) and cLIR (dark) indicated along the sequence. FFAT, two phenylalanines in an acidic tract; Pro, proline-rich motif; RBD, Rab11-binding domain; pLDDT, predicted local distance difference test; ipTM, interface predicted template modeling; PAE, predicted aligned error; LIR, local interaction residue (PAE ≤ 12 Å); cLIR, contact-filtered LIR (PAE ≤ 12 Å and Cβ ≤ 8 Å).

Predicted template modeling (pTM) and predicted local distance difference test (pLDDT) scores are lowered by large disordered regions such as the WDR44 N-terminus even where a local interface is confidently predicted. Predicted aligned error (PAE), which estimates the error associated with the relative positions of two residues, does not share the same limitation. To map residues at inhibitory contacts, we used the PAE-based Local Interaction Visualization and Analysis (LIVIA) platform (Kim and Perrimon, 2026). LIVIA designates a residue a local interaction residue (LIR) when the model places it confidently relative to a residue in the partner chain, and a contact-filtered LIR (cLIR) when that pair is also close enough to be in direct contact. This analysis identified two discrete interfaces: a ‘top-binding’ segment (383-407) in the N-terminus that is confidently predicted to associate with the “top” surface of the C-terminal WD40 domain, and a ‘side-binding’ segment (301-321) that contacts the side of the WD40 domain (Figure 2, B-D; Figure S1B). Using PISA analysis (Krissinel and Henrick, 2007), we found the buried surface area (BSA) at the top and side interfaces to be extensive, totaling 920 Å^2^ and 1119 Å^2^, respectively (Table S1).

AlphaFold3 modeling (Abramson *et al*. 2024) and LIVIA analysis determined that the predicted side-binding and top-binding interfaces are structurally conserved across WDR44 orthologs in humans, frogs, flies, and fission yeast, further supporting their functional importance (Figure S1C), despite the absence of a Rab11-binding domain in yeast (Figure S1D). Taken together, these in silico analyses identify two regions of the WDR44 N-terminus that form extensive interfaces with distinct surfaces of the WD40 domain.

### Both interfaces are required to maintain the closed conformation

To assess whether AlphaFold2-predicted contacts are responsible for maintaining the ‘closed’ conformation of the protein, we designed mutations to disrupt each intramolecular interface. The top-binding segment contains a helix that is confidently predicted to contact the top face of the WD40 domain, as previously reported (Thibodeau et al., 2023), and lies close to disease mutation sites D648G, L668S, and D669N (Figure 3A). To displace the helix, we mutated a threonine at its base to glutamic acid, introducing both steric hindrance and negative charge (WDR44^T385E^; Figure 3A). The interface with the side-binding segment was mutated by deleting residues 301-321 (WDR44^Δ301-321^). To determine if loss of these intramolecular contacts promotes Rab11 binding, we used BRET to detect endosomal localization of single and double interface mutants (Figure 3B). Single mutants showed increased recruitment to Rab11-positive compartments, confirming these mutations disrupt autoinhibitory regions. Displacing both contacts had an even stronger effect, demonstrating that both interfaces contribute to autoinhibition.

**Figure 3.**
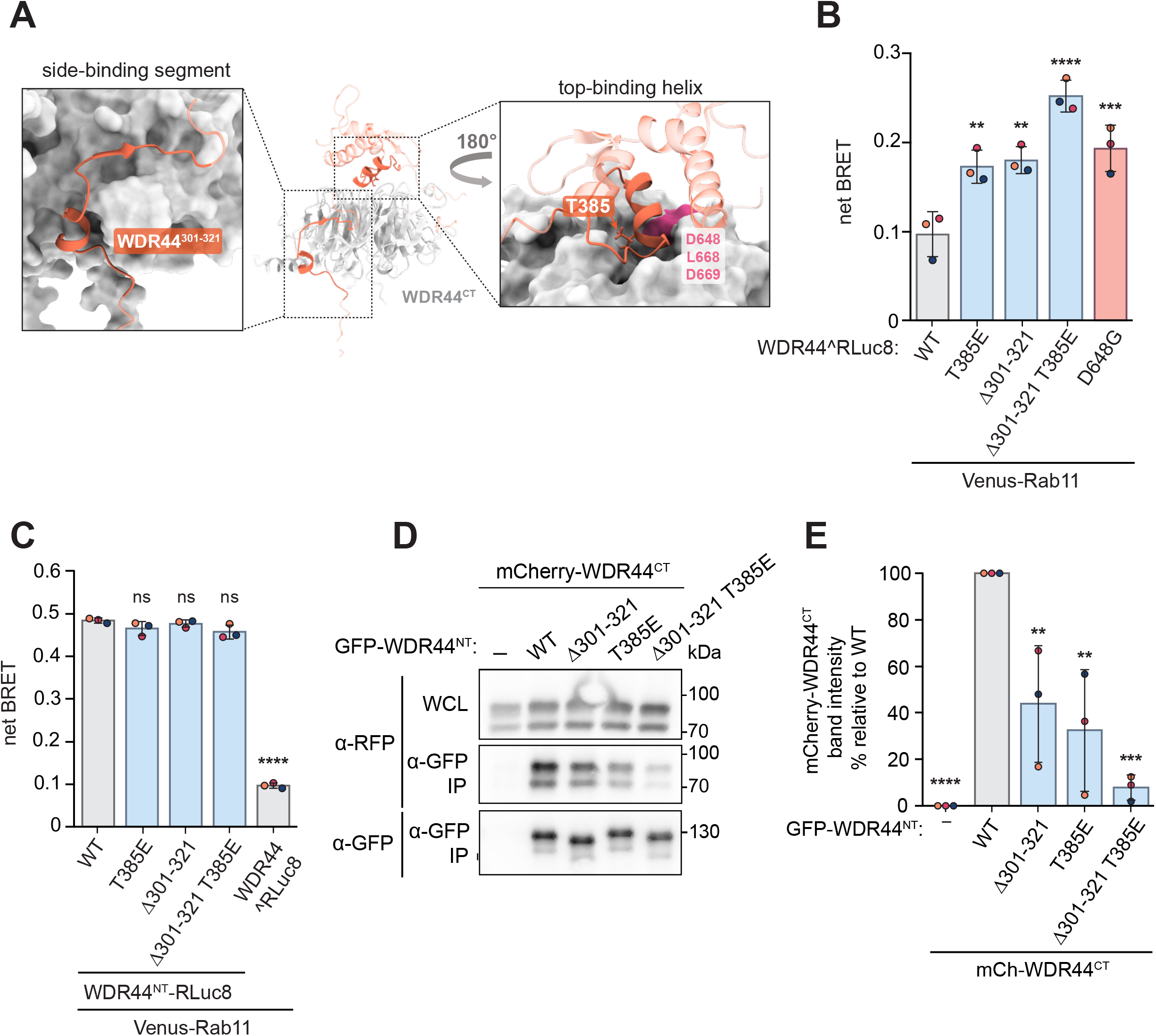
Displacement of both intramolecular interfaces relieves WDR44 autoinhibition. **(A)** AlphaFold2-predicted structure of the N-terminal side-binding segment and top-binding helix (orange) with the C-terminus (grey). Insets highlight the regions selected for mutation: T385 in the top-binding helix, and residues 301-321 in the side-binding segment. Three pathogenic mutation sites proximal to the top-binding helix are indicated (magenta). **(B)** Displacement of the side-binding segment or top-binding helix causes WDR44 re-distribution to Rab11-positive compartments, quantified by BRET analysis. The pathogenic variant D648G is included as a benchmark, and all comparisons are to WT. One-way ANOVA with Dunnett’s multiple comparison’s test; n = 3; ** = P ≤ 0.01, *** = P ≤ 0.001, **** = P < 0.0001. Error bars indicate standard deviation. **(C)** Localization of the WDR44 N-terminus, C-terminally tagged with RLuc8, is unaffected by mutations in the side- and top-binding segments, quantified by BRET analysis. Full-length WDR44 is shown for comparison. One-way ANOVA with Dunnett’s multiple comparison’s test; n = 3; ns = P > 0.05, **** = P < 0.0001. Error bars indicate standard deviation. **(D)** Co-immunoprecipitation of transiently transfected mCherry-WDR44^CT^ with GFP-WDR44^NT^ is disrupted by mutation of the side-binding segment or top-binding helix. GFP-tagged proteins were immunoprecipitated from HeLa cells and probed for GFP and RFP. A representative blot of three independent experiments is shown; the first lane received no GFP-tagged construct. **(E)** Quantification of the co-IP of mCherry-WDR44^CT^ with GFP-WDR44^NT^ mutants in D by densitometry. All values are relative to the WT protein. One-way ANOVA with Dunnett’s multiple comparison’s test; n = 3; ** = P ≤ 0.01, *** = P ≤ 0.001, **** = P < 0.0001. Error bars indicate standard deviation. BRET, bioluminescence resonance energy transfer; WCL, whole cell lysate; IP, immunoprecipitate; co-IP, co-immunoprecipitation; GFP, green fluorescent protein; RFP, red fluorescent protein.

Mutations in the WDR44 N-terminus could alter the structure of the Rab11-binding site to enhance Rab11 binding, independent of intramolecular interactions involving the C-terminal WD40 domain. To address this, we tested if these same mutations affect recruitment of the isolated WDR44 N-terminal fragment, C-terminally tagged with RLuc8, to Rab11 compartments (Figure 3C). We confirmed that BRET strongly detected the isolated N-terminus at Rab11-labeled compartments and this was unchanged by mutation of the top-binding and side-binding segments. This supports the conclusion that mutations in top and side-binding regions disrupt autoinhibitory contacts.

BRET reports WDR44 recruitment to Rab11 compartments, which provides a readout of the open conformation, but does not directly demonstrate contacts between the N- and C-termini. To test if these mutations disrupt interactions between the WDR44 N- and C-termini, we performed co-immunoprecipitation (co-IP) of GFP-WDR44^NT^ and mCherry-WDR44^CT^ fragments. Mutations predicted to displace the top-binding helix or side-binding segment individually decreased binding and resulted in a strongly reduced interaction when combined (Figure 3, D and E), indicating both sites contribute to the intramolecular interaction. Collectively, our data define two internal interfaces that regulate WDR44-Rab11 binding by maintaining the autoinhibited conformation.

### An unstructured strand extends the top-binding interface

The top-binding region of the N-terminus, defined by our LIVIA analysis, contains an unstructured strand that is less confidently modeled than the helical region, yet is predicted to contact the C-terminal WD40 domain close to two pathogenic mutation sites, S764F and G782C (Figure 4A). To test whether this region contributes to autoinhibition, we created a charge-swap mutant by replacing four negatively charged residues with lysine (WDR44^4K^) (Figure 4, A and B). This mutant showed increased endosomal localization that was comparable to that of a mutant containing the S764F pathogenic variant (Figure 4C). A second mutant that instead truncates six strand side chains to alanine (WDR44 6A) similarly showed that disruption of the strand region alone was sufficient to relieve autoinhibition (Figure 4, A-C). Thus, the strand is an important component of the top-binding interface that maintains the closed conformation of WDR44.

**Figure 4.**
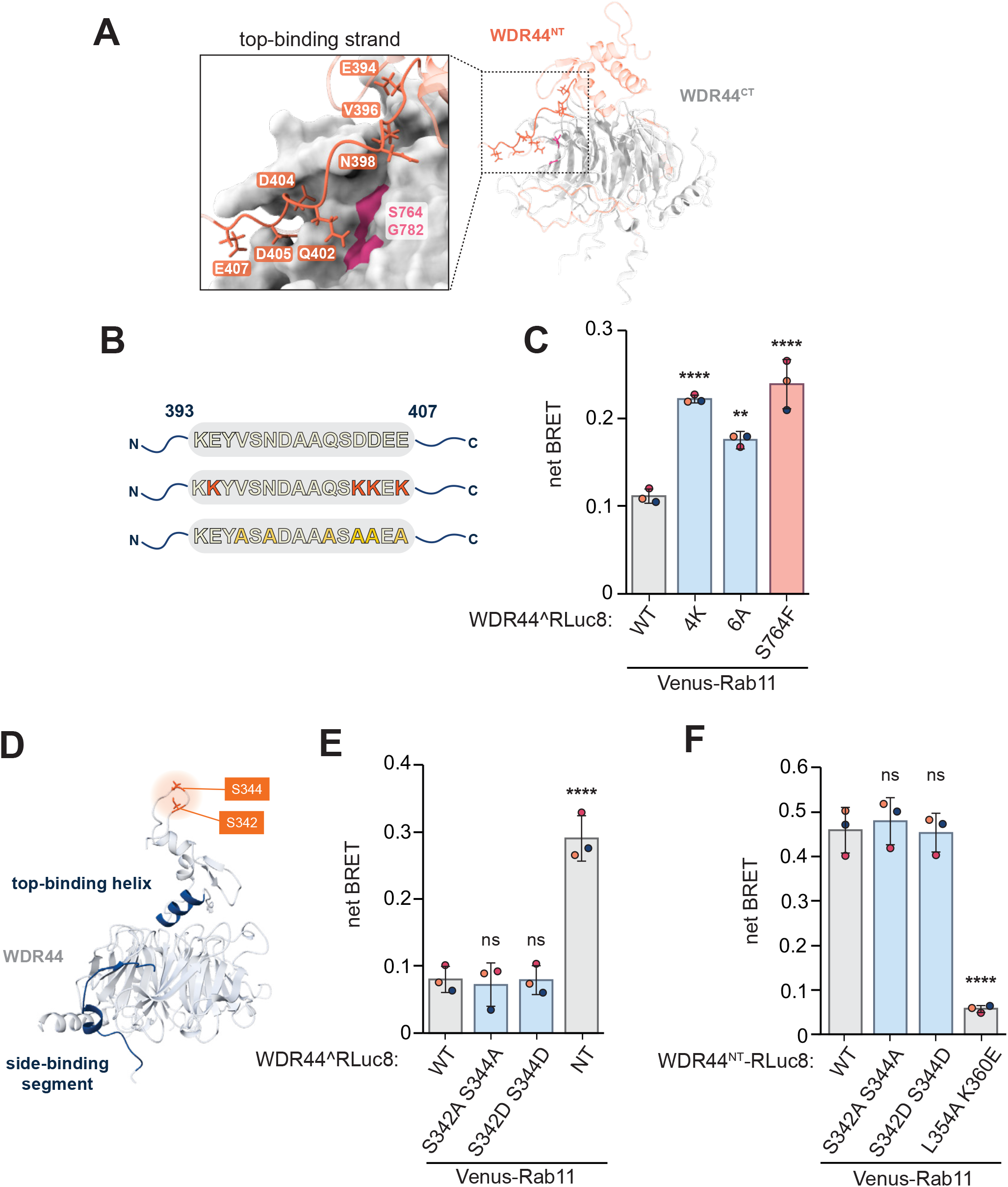
An unstructured strand in the top-binding interface contributes to autoinhibition. **(A)** AlphaFold2-predicted structure of the WDR44 showing the N-terminal top-binding strand (orange) and C-terminus (grey). Inset highlights mutated residues in the strand, which are proximal to two pathogenic mutation sites in the WD40 domain (magenta). **(B)** Schematic depicting the sequence of the strand, highlighting the four negatively charged residues mutated to lysine (red), and the six residues mutated to alanine (yellow). **(C)** Displacement of the top-binding strand induces Rab11 proximity of WDR44, measured by BRET. The pathogenic variant S764F is included for comparison. One-way ANOVA with Dunnett’s multiple comparison’s test; n = 3; ** = P ≤ 0.01, **** = P < 0.0001. Error bars indicate standard deviation. **(D)** AlphaFold2-predicted structure of WDR44, highlighting known phosphorylation sites (orange) that are not located within autoinhibitory regions of the N-terminus (blue). The highest ranked model is shown and residues with pLDDT values < 30 are hidden. **(E)** Localization of full-length WDR44 at Rab11-labeled structures is unaffected by phosphomutations at S342 and S344, quantified by BRET analysis; the isolated N-terminus (NT), internally tagged with RLuc8 at position 292, is shown as a positive control. One-way ANOVA with Dunnett’s multiple comparison’s test; n = 3, ns = P > 0.05, **** = P < 0.0001. Error bars indicate standard deviation. **(F)** Rab11 proximity of the WDR44 N-terminal fragment, C-terminally tagged with RLuc8, is unchanged by mutations at S342 and S344, while point mutations in the Rab11-binding domain cause a loss of recruitment. One-way ANOVA with Dunnett’s multiple comparison’s test; n = 3; ns = P > 0.05, **** = P < 0.0001. Error bars indicate standard deviation. BRET, bioluminescence resonance energy transfer; pLDDT, predicted local distance difference test; NT, N-terminus.

Phosphorylation of residues S342 and S344 by Akt and Sgk3 is reported to promote WDR44-Rab11 interactions (Walia *et al*., 2019, Malik *et al*., 2019, Thibodeau *et al*., 2023). These residues lie within the Rab11 binding domain, yet are not predicted to directly contact Rab11 (Figure 4D; Thibodeau *et al*., 2023), nor are they close to our modeled autoinhibitory contacts. To understand how phosphorylation regulates WDR44-Rab11 interactions, we created S342 S344 phosphomimetic or phosphomutants previously reported to impact Rab11 binding (Walia *et al*., 2019). While the internally RLuc8-tagged WDR44 N-terminus showed recruitment to Rab11-positive membranes, comparable to that of mutants that promote an open conformation (Figure 3B), mutations at S342 and S344 did not alter Rab11 proximity, suggesting modification of these residues does not strongly induce the open conformation (Figure 4E).

To test if phosphorylation of these residues affects the Rab11-binding interface directly, we used BRET to measure localization of the isolated WDR44 N-terminal fragment tagged at the C-terminus with RLuc8 (Figure 4F). As expected, an L354A and K360E double mutant, which contains two point mutations previously shown to block WDR44-Rab11 binding *in vitro* (Thibodeau *et al*., 2023), strongly reduced Rab11 recruitment. In contrast, neither phosphomimetic nor phosphodeficient S342 S344 mutants showed altered Rab11 proximity. While the strong BRET signal may obscure small changes in Rab11 binding, these results show that mutation of these sites does not abolish the WDR44-Rab11 interface. This is consistent with a previous study (Walia *et al*., 2019), which found alanine substitutions at S342 and S344 reduce but do not block Rab11 binding, and suggested that phosphorylation at these sites may not be sufficient to explain the inhibitory effects of WDR44 on ciliogenesis. Two sites in the WDR44 top-binding strand, S397 and S403, are phosphorylated in phorbol myristate acetate (PMA)-treated cells (Galan *et al*., 2014), but phosphomimetic and phosphodeficient substitutions at these sites also had no effect (Figure S2, A and B). Future studies will be needed to determine whether other WDR44 phosphorylation events could regulate Rab11 binding by affecting autoinhibitory contacts.

Together, this work defines two main regions in the WDR44 N-terminus that form intramolecular interactions with the C-terminus. The top-binding segment extends along the surface of the WD40 domain, and encompasses five of the seven reported pathogenic point mutation sites. Although H839R, which maps outside these autoinhibitory interfaces, could act indirectly by perturbing the overall folding of the WD40 domain (Accogli et al., 2024), the adjacent N840S mutation did not increase Rab11 proximity in our assay and may instead be pathogenic through a mechanism unrelated to autoinhibition. In contrast, no pathogenic variants have yet been mapped to the side interface. It is tempting to speculate that this interface has a regulatory role that makes it intolerant to mutation. In fact, evolutionary conservation of both the top and side interfaces, even in organisms such as *S. pombe*, whose WDR44 ortholog lacks a recognizable Rab11 binding domain, suggests that these autoinhibitory contacts may be important for Rab11-independent functions of WDR44.

WDR44 is a suppressor of ciliogenesis, but only in serum-fed conditions (Parolek and Burd, 2024). Current models suggest that phosphorylated forms of WDR44 sequester Rab11 and prevent its interaction with the ciliogenesis-promoting FIP3-Rabin8 complex. WDR44 is also required for cargo traffic at tubular recycling endosomes. This could indirectly affect ciliogenesis initiation, given that preciliary vesicles are derived from this compartment (Saha et al., 2024) and that WDR44-positive endosomes are enriched with early ciliogenesis proteins including EHD1, MICAL1, and Rab8 (Lucken-Ardjomande Häsler *et al*., 2020, Parolek and Burd, 2024). If WDR44 promotes cell surface-directed trafficking of endosomal cargo, it may counteract the formation or delivery of preciliary vesicles, though this hypothesis remains unexplored. Further studies are needed to determine if the inhibitory interfaces we have identified contribute to the regulation of ciliogenesis, and to understand how WDR44 autoinhibition regulates its downstream functions.

## Materials and Methods

### Plasmids

Plasmid information and primer sequences used in this study are listed in Tables S2 and S3. Venus-tagged Rab11, Rab5, and Rab7 were gifts from Dr. Nevin Lambert (Augusta University, Augusta, GA, USA; Lan et al., 2012). Plasmids expressing mutant WDR44 were constructed using Golden Gate cloning in *Escherichia coli*.

### Structural modeling and analysis

Structural models of proteins and binding interfaces were generated using AlphaFold2-powered ColabFold (Jumper et al., 2021; Mirdita et al., 2022) and the AlphaFold3 server (Abramson *et al*. 2024). LIVIA (Kim e*t al.* 2024, Kim and Perrimon, 2026, Kim *et al*., 2026) was used to compute local protein-protein interaction metrics. Local interaction residues (LIR) were defined as inter-chain residue pairs with a predicted aligned error of 12 Å or less, and contact-filtered LIR (cLIR) as the subset whose Cβ atoms lie within 8 Å. Interface analysis was performed using the PDBePISA server (Krissinel and Henrick, 2007) on the highest ranked model. Structural models were generated for presentation on UCSF ChimeraX v1.12 (Pettersen et al., 2021).

### Cell culture and transfection

HEK293T and HeLa cells from ATCC were grown in high-glucose Dulbecco’s Modified Eagle Medium (DMEM; Gibco) supplemented with 10% fetal bovine serum (FBS; Gibco), 4 mM L-glutamine, and 1 mM sodium pyruvate in a humidified incubator at 37°C, 5% CO_2_. For BRET assays, HEK293T cells were grown on 6-well plates to 80% confluency and transfected with 0.05 μg of donor construct DNA and 0.5 μg of acceptor construct DNA using Lipofectamine 2000 (Invitrogen) as per manufacturer’s instructions. For immunoprecipitation, HeLa cells were grown on 60 mm plates to 80% confluency and transfected with 1 μg of each DNA.

### BRET assays

24 hours after transfection, cells were harvested using 0.25% trypsin-EDTA (Gibco), washed with PBS, and replated on Nunc F96 MicroWell White Polystyrene 96-well plates (Thermo Fisher Scientific). Coelenterazine h (5 μM; Cayman) was added to cells. A Tecan Spark multimode microplate reader (Tecan) was used to inject the substrate and immediately obtain fluorescence and luminescence measurements at 485 and 530 nm. To calculate raw BRET signals, the acceptor emission (530 nm) was divided by the donor emission (485 nm). Net BRET values were calculated as the difference between the raw BRET ratios of cells co-expressing donor and acceptor minus raw BRET ratios of cells expressing the donor only.

### Immunoprecipitation and western blotting

Cells were washed with PBS and harvested 24 hours after transfection using 0.25% trypsin-EDTA (Gibco). Cells were washed again with PBS before lysis in 300 μl lysis buffer (150mM NaCl, 1mM EDTA, 20mM Tris-Cl pH 8.0, 1% TritonX-100). Protein levels were measured using a BCA assay kit (Thermo Fisher Scientific). A fraction of protein lysate was resuspended in 2x Laemmli sample buffer (240 mM Tris-Cl pH 6.8, 8% SDS, 40% glycerol, 0.004% bromophenol blue, 1% β-mercaptoethanol). The remainder of protein lysate was incubated with polyclonal rabbit anti-GFP (EU2; Eusera) for 1 h, followed by Protein-A-Sepharose beads (GE Healthcare) for 1 h, all at 4°C. Beads were washed and eluted in 50 μl Thorner buffer (40 mM Tris pH 6.8, 8 M urea, 5% SDS, 0.1 M EDTA, 0.4 mg/ml bromophenol blue, 1% β-mercaptoethanol) for 5 min at 70°C.

Lysate and immunoprecipitated samples were separated on 7% SDS-PAGE gels, followed by transfer to nitrocellulose membranes. Membranes were blotted with mouse monoclonal anti-GFP (11814460001; Roche) or mouse monoclonal anti-RFP (6G6; Chromotek) primary antibodies then horseradish peroxidase–conjugated polyclonal goat anti-mouse (115-035-146; Jackson ImmunoResearch Laboratories). Blots were developed with ECL chemiluminescent reagents (Cytiva) and imaged using the Vilber Fusion FX. Densitometry was performed using Fiji.

### Statistical analysis

GraphPad Prism 11 (GraphPad Software, San Diego, California) was used to conduct statistical analyses as described in figure legends. All n values refer to independent biological replicates performed on separate days; bars show the mean and error bars the standard deviation. Normal data distribution was assumed but not formally tested.

## Data Availability Statement

All data generated in this study are included in the manuscript and supplemental files.

## Acknowledgements

We thank Dr. Steve Caplan (University of Nebraska Medical Center) for insightful discussions. This work was supported by the Canadian Institutes of Health Research (grants OGB-177941 and PJT-180544).

## Abbreviations

BRET: bioluminescence resonance energy transfer
LIR: local interaction residue

---

**Figure S1.**
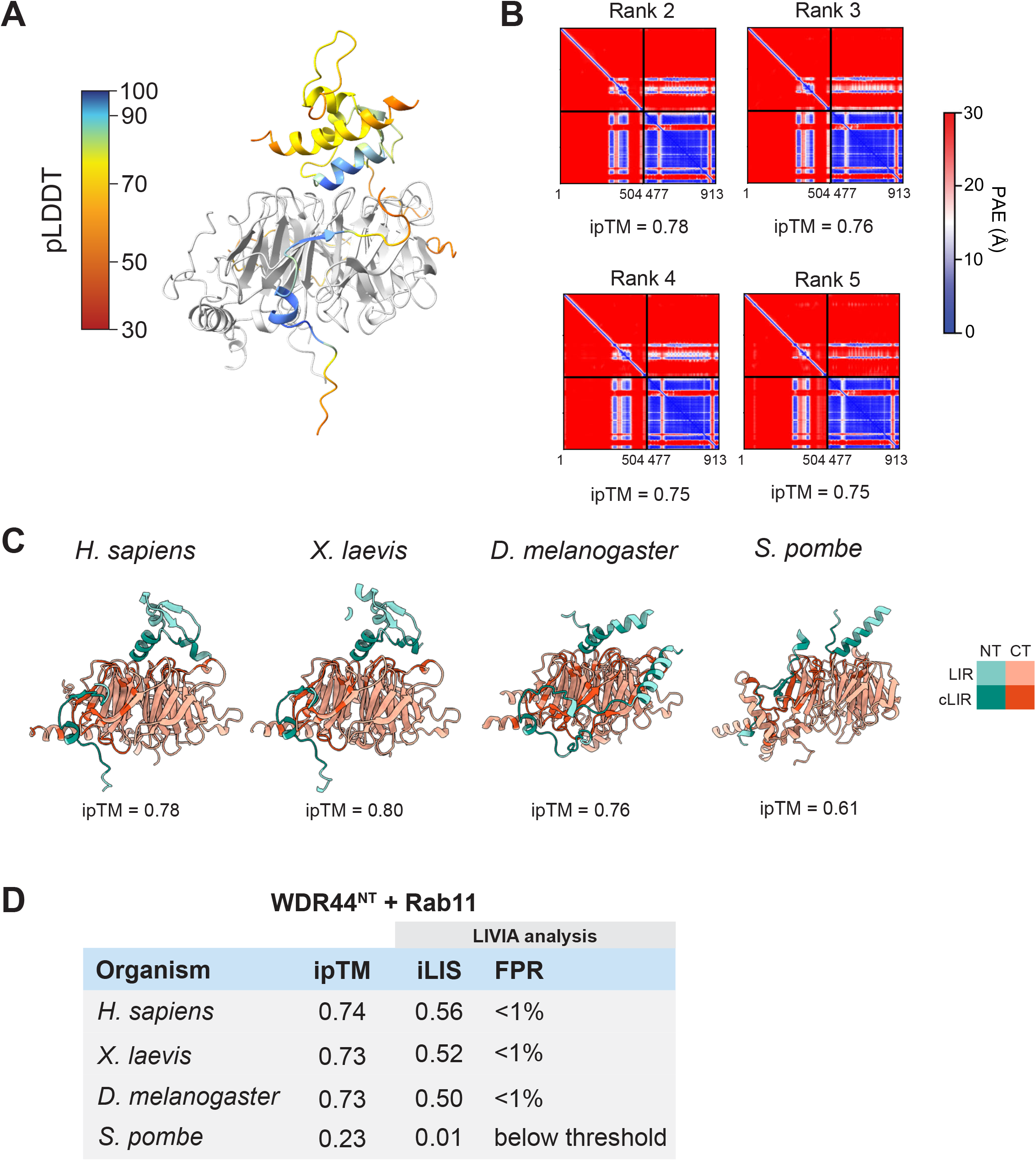
Model confidence and cross-species conservation of the WDR44 intramolecular interfaces**. (A**) AlphaFold2-predicted structure of the WDR44 N-terminus, colored by per residue pLDDT values and the C-terminus (grey). The highest ranked model is shown and residues with pLDDT values < 30 are hidden. **(B)** PAE plots for the AlphaFold2-based predictions shown in Figure 2A of models ranked 2-5. ipTMs of each ranked model are indicated. **(C)** AlphaFold3-predicted interaction between the WDR44 N- (green) and C- (orange) termini in *H. sapiens, X. laevis, D. melanogaster, and S. pombe* orthologs. LIVIA-computed local interaction residues (LIR, light green/orange) and the subset also in direct contact (cLIR, dark green/orange) mark the highest confidence interfaces. (D) LIVIA-computed confidence metrics for the AlphaFold3-modeled interaction between the WDR44 N-terminus and Rab11 in *H. sapiens*, *X. laevis*, *D. melanogaster*, and *S. pombe* orthologs. ipTM scores of the highest ranked model are indicated. Average iLIS scores of five models, and the resulting FPRs are indicated. pLDDT, predicted local distance difference test; PAE, predicted aligned error; LIR, local interaction residue (PAE ≤ 12 Å); cLIR, contact-filtered LIR (PAE ≤ 12 Å and Cβ ≤ 8 Å); ipTM, interface predicted template modeling; iLIS, integrated local interaction score; FPR, false positive rate.

**Figure S2.**
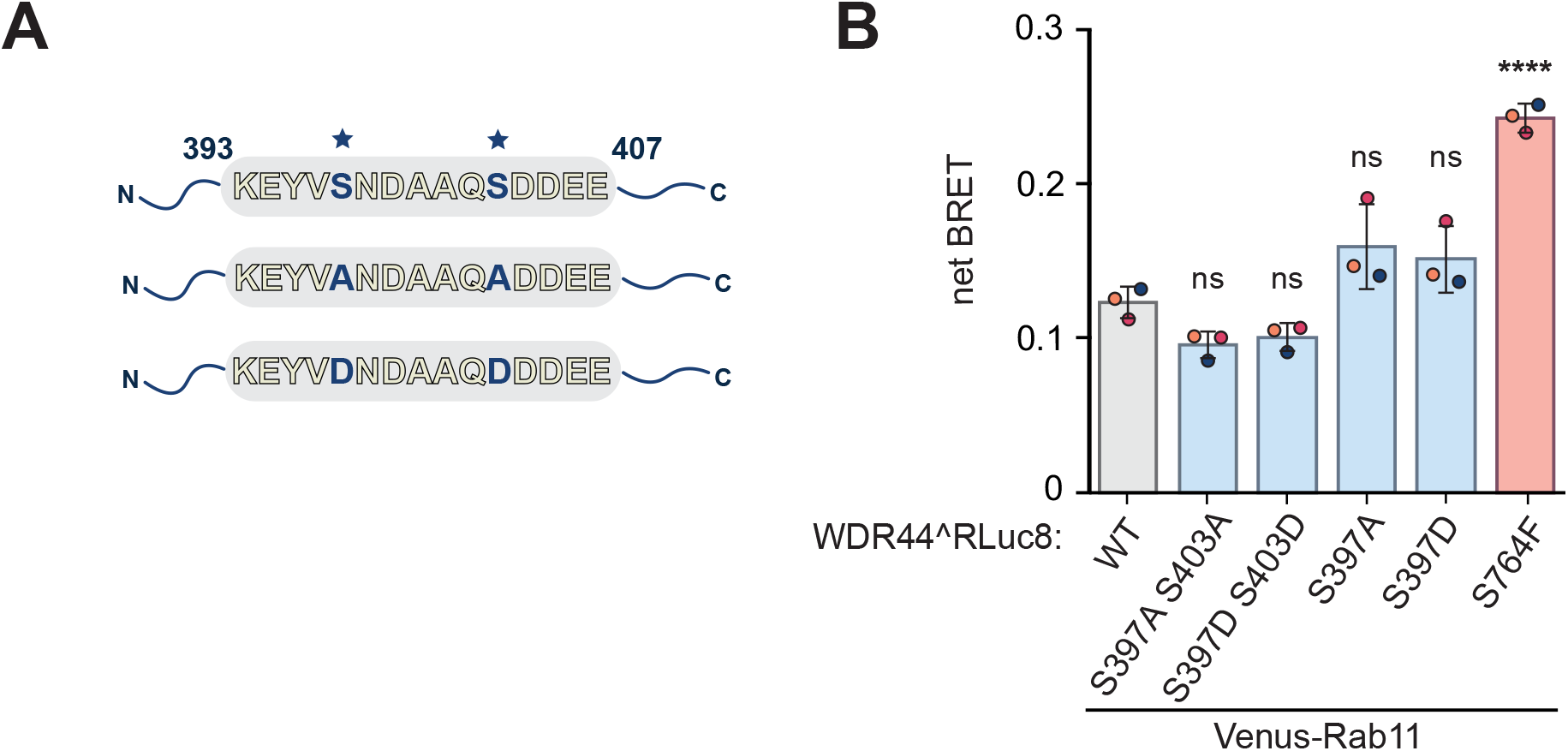
Phosphomutations at S397 and S403 do not alter WDR44 recruitment**. (A)** Schematic depicting the sequence of the unstructured strand, highlighting two phosphorylated serines mutated to alanine or aspartic acid (blue). **(B)** Single and double phosphomimetic and phosphodeficient substitutions at S397 and S403 do not alter WDR44 localization at Rab11-labeled compartments, quantified by BRET analysis. The pathogenic variant S764F is included as a positive control. One-way ANOVA with Dunnett’s multiple comparison’s test; n = 3, ns = P > 0.05, **** = P < 0.0001. Error bars indicate standard deviation. BRET, bioluminescence resonance energy transfer.

